# Population genetics meets precision-cut kidney slices: Nephrotoxicity modelled ex vivo in the founder strains of the BXD mouse consortium

**DOI:** 10.64898/2026.06.16.732581

**Authors:** Jana Andres, Nichakorn Phengpol, Mikhail Burmakin, Hannes Olauson, Jaakko Patrakka, Matthias B. Moor

## Abstract

Acute kidney injury (AKI) affects millions of patients annually and is associated with high morbidity and mortality, to date no curative treatment exists. Drug-induced nephrotoxicity accounts for up to 25% of AKI cases, but individual susceptibility remains hard to predict. While genetic factors are suspected to play a part in this variability, the pharmacogenomics of nephrotoxin-induced kidney injury remain largely unknown.

To investigate genetically determined susceptibility, we used precision-cut kidney slices (PCKS) from the two founder strains of the BXD mouse consortium, C57BL/6J and DBA/2J. PCKS preserves tissue architecture and cell-cell interaction, allowing close experimental control while maintaining the renal microenvironment. Slices were exposed to cyclosporine A (80 nM for 6h, 20nM for 24h and 48h) and Tunicamycin (1 µM for 6h and for 24h) as well as normoxia (4°C for 20h) and hyperoxia (4°C for 20h and 4h in incubator). Slices were then analysed using histopathological scoring, TUNEL staining, ATP quantification and bulk RNA sequencing.

We found that the main source of variation was experimental duration. Nevertheless, a subtle difference between the strains could be observed for both cyclosporine A and Tunicamycin, with DBA/2J showing a stronger response to nephrotoxic stress, including lower ATP levels, higher proportion of apoptotic cells and a more pronounced transcriptomic response. For both strains, normoxia was the least harmful condition.

These findings support the hypothesis that the BXD founder strains differ in their susceptibility to nephrotoxic kidney injury and support the use of PCKS as a relevant ex vivo model for studying early renal stress response. This provides the foundation to extend this approach to a broader spectrum of the BXD population to identify genetic loci and candidate genes involved in genetic susceptibility to nephrotoxins.

## Introduction

Acute kidney injury (AKI) remains a clinical challenge worldwide, affecting up to 13.3 million individuals per year.^1^ Approximately 7% of hospitalized patients develop AKI, the majority of whom are admitted for non-renal conditions.^2^ To date there is no existing cure, management is largely limited to supportive measures.^3^ Although AKI is rarely the direct cause of death, it is associated with increased morbidity, mortality, and significant healthcare cost. Mortality is especially high in patients requiring dialysis.^2,4,5^ While most patients experience a complete or partial recovery, AKI confers a risk of progression to end-stage renal disease (ESRD), particularly among patients with preexisting renal impairment or in older patients.^6–8^ Patient populations are increasingly older and present with a higher burden of comorbidities. They present with more complex clinical conditions and are consequently exposed to an increasing number of potential pharmacological treatments, all of which may contribute to a higher risk of developing AKI.^9^

Drug-induced nephrotoxicity accounts for up to 25 percent of AKI cases in the general population (including both in- and outpatient settings)^2,5,9,10^ and up to 66 percent in elderly patients.^11^ It therefore presents one of the leading causes of AKI, frequently limits drug dosage or therapeutic options, thereby negatively affecting patient outcomes. Although several mechanisms of drug-induced nephrotoxicity, including acute tubular injury, tubulointerstitial inflammation and crystal formation of insoluble compounds, are well described, it remains unclear why some patients develop AKI while others do not.^3,10,11^ This may indicate the presence of an individual, genetically determined susceptibility to drug-induced nephrotoxicity.

A genome-wide association study of patients receiving platinum-based chemotherapy has identified several genetic variants associated with AKI,^12^ suggesting that pharmacogenomic approaches could potentially be used to help predict susceptibility to treatment-induced nephrotoxicity. However, genetic predisposition to nephrotoxicity from other commonly used drugs, including antibiotics, non-steroidal anti-inflammatory drugs, and calcineurin inhibitors, remains largely unknown.

To investigate the contribution of genetically determined susceptibility, well-characterized genetic reference populations (GRPs) are required. In mammals one of the best studied and largest GRP is the BXD mouse family.^13,14^ The BXD strains originate from a cross between the two inbred parental lines, C57BL/6J and DBA/2J, followed by successive inbreeding over multiple generations, resulting in more than 150 recombinant, inbred, isogenic lines, that serve as an immortalized genetic mapping population.^15^ Because the founder strains of the BXD-family are fully inbred and have a well-characterized genome,^16,17^ the resulting BXD lines have fixed genotypes. This enables precise investigation of genotype-phenotype relationships across strains and close experimental control.

Previous studies have successfully used the BXD-consortium to identify quantitative trait loci (QTLs) and underlying genes associated with complex traits and disease models across a wide range of phenotypes, including metabolic disorders, cardiovascular disease and bone density.^13,14,18–21^

Although animal models have been used to effectively model and study AKI, including pathological mechanisms and potential biomarkers, they have shown limited ability to predict clinical efficacy of potential therapeutic agents. In vitro models provide valuable insights into pathogenic mechanisms, however they do not reflect the complexity of the in vivo milieu and lack cellular heterogenity.^4,22–25^

Recent studies have successfully used Precision-cut kidney slices (PCKS) derived from different species to model various forms of kidney injury, including endoplasmic reticulum stress, ischemia-reperfusion injury following transplantation, fibrosis and anti-fibrotic drug development.^22,23,26,27^ PCKS offers a multicellular ex vivo model that preserves tissue architecture and cell-cell interactions and may therefore represent a valuable alternative to existing, limited models.^28^ In addition, their use contributes to the reduction, refinement and replacement principle (“3Rs” principles) of animal studies.^29^ Another major advantage of PCKS, compared with classical models, is the applicability to human kidney tissue, enabling functional studies that are otherwise largely limited to descriptive analyses.^22^

In this study, we aim to investigate the hypothesis that the ancestral strains of the BXD mouse consortium exhibit differences in their susceptibility to nephrotoxin-induced kidney injury. To test this hypothesis, we used PCKS from C57BL/6J and DBA/2J mice to compare several nephrotoxicity-related phenotypes. Specifical we will evaluate histopathological tissue damage to assess structural alterations indicative for kidney injury, measure ATP levels as a key indicator of cellular metabolic activity, quantify apoptosis to understand the extent of cell death and perform bulk RNA sequencing to analyse transcriptomic changes that underlie responses to nephrotoxin exposure.

Demonstrating phenotypic variation among these ancestral strains would justify the use of this genetically diverse BXD F2-derived recombinant inbred resource for identifying genetic modifiers of susceptibility to nephrotoxin-mediated kidney injury through genome-wide association studies (GWAS) or QTL-mapping. Ultimately this approach could lead to the identification of causal genetic variants associated with nephrotoxin-induced phenotypes, such as renal tubular cell apoptosis or biochemical signs of impaired tissue viability.

## Materials and methods

### Animal Experiments

Male C57BL/6J and DBA/2J mice aged 8–10 weeks were obtained from Charles River and utilized in this study. A total of six mice per strain were used. All animal procedures were conducted in accordance with the guidelines of the German Society for Laboratory Animal Science (Gesellschaft für Versuchstierkunde, GV-SOLAS). Animals were housed in standard individually ventilated cages under strictly controlled environmental conditions, including a 12 h light/dark cycle, ambient temperature of 20 ± 2 °C, and relative humidity of 50 ± 5%. Standard laboratory chow and water were provided ad libitum throughout the experimental period. Upon arrival, mice were randomly assigned to experimental groups using body weight–based stratification to ensure comparable baseline characteristics across groups. procedures were carried out at the Preclinical Laboratory, Karolinska Institutet, and were approved by the Linköping Animal Experimentation Ethics Committee under ethical approval number DNR 19667-2022.

### PCKS and pertubation protocols

After decapsulation, kidney tissues were embedded in 2% agarose warmed to 40 °C. Tissue sections of 300 µm thickness were then prepared using a vibrating microtome (Compresstome® VF310-0Z, PrecisionaryInstruments) in ice-cold DPBS. Slices were promptly transferred into Williams’ Medium E supplemented with GlutaMAX™-I (Gibco™, Thermo Fisher Scientific), 25 mM D-glucose, and 1% penicillin-streptomycin (10,000 U/mL, Gibco™, Thermo Fisher Scientific). To induce nephrotoxicity, kidney slices were exposed to cyclosporine A (20 and 80 µM; 30024, Sigma-Aldrich) or tunicamycin (0.1, 0.5, and 1 µM; T7765, Sigma-Aldrich; #3516, Tocris Bioscience) for 6, 24, and 48 h. In the normoxia–hyperoxia model, tissue slices were maintained at 4°C for 20 h, followed by incubation for an additional 4 h prior to harvesting. Tissue samples were collected and preserved either in 10% formalin (HT501128, Sigma-Aldrich) for histological analysis or in RNAlater for subsequent RNA extraction. Samples designated for ATP analysis were snap-frozen and stored at −80°C. Each experimental condition was performed in triplicate (technical replicates).

### H&E Staining

Kidneys were fixed with 4% paraformaldehyde, subsequently embedded with paraffin, sectioned at 4µm. Hematoxylin and eosin (H&E) staining was performed using standard laboratory procedures. Renal injury was evaluated by an observer blinded to group allocation using a validated semiquantitative scoring system (0–4) based on the percentage of cortical area affected (0 = none; 1 = <25%; 2 = 25–50%; 3 = 51–75%; 4 = >75%), following criteria: small basal vacuoles, larger non-basal vacuoles, cell detachment, denuded basal membranes, loss of nuclei, and autolysis. Six randomly selected, non-overlapping cortical fields per group (8× magnification) were evaluated and averaged to generate a single composite score per sample.

### TUNEL Staining

TUNEL staining was performed to assess apoptotic DNA fragmentation in paraffin-embedded kidney sections using an HRP-DAB detection system, according to the manufacturer’s protocol with minor procedural modifications. Briefly, sections were deparaffinized, rehydrated through a graded ethanol series, rinsed with TBS, and maintained in a hydrated state throughout the staining procedure. Tissue sections were permeabilized with Proteinase K, and endogenous peroxidase activity was quenched prior to TdT-mediated nucleotide labelling. Following the labelling reaction, sections were incubated with HRP conjugate and developed with DAB chromogen, producing dark brown nuclear staining in apoptotic cells. The slides were counterstained with methyl green, dehydrated, cleared, mounted, and examined under a light microscope. Apoptotic cells were identified based on DAB-positive nuclear staining together with characteristic morphological features.

### Quantifying TUNEL Stanning

For the analysis of the TUNEL-stained PCKS, the open source software QuPath (version 0.6.0) was used.^30^ To ensure accurate automatic cell detection, cell-detection parameters were established by a single investigator who was blinded to the treatment groups at the time of analysis. The parameters were calibrated by manually annotating a representative tissue section containing approximately 4500 cells (in sample 2A-1). The automatic cell detection was then optimized to match the manual counts, using the following parameters: threshold of 0.38, pixel size of 0.3, intensity threshold of 0.271. A maximum area threshold of 33µm^2 was added to exclude oversized, non-nuclear, background-staining artefacts. Based on these settings the nuclei were then automatically classified as either TUNEL-positive or TUNEL-negative.

### Tissue ATP quantification

ATP levels were quantified using an ATP Assay Kit (Colorimetric/Fluorometric; ab83355, Abcam) according to the manufacturer’s protocol. Frozen kidney slices, approximately 3 mg of tissue per sample, were homogenized in 30 µL of ATP assay buffer. The homogenates were then incubated with 50 µL of ATP reaction mix for 30 min at room temperature in the dark. Absorbance was measured at 570 nm. ATP concentrations were calculated from a standard curve and normalized to total protein concentration, which was determined using the Pierce™ BCA Protein Assay Kit.

### RNA extraction and sequencing

Two pooled kidney slices per condition were minced in RLT buffer + DTT. RNA was extracted using QIAGEN RNeasy Mini Kit (QIAGEN) according to manufacturer’s instructions. Total RNA was subjected to quality control with Agilent Tapestation according to the manufacturer’s instructions. To construct libraries suitable for Illumina sequencing, the Illumina stranded mRNA prep ligation sample preparation protocol was used with starting concentration between 25-1000 ng total RNA. The protocol includes mRNA isolation, cDNA synthesis, ligation of adapters and amplification of indexed libraries. The yield and quality of the amplified libraries was analyzed using Qubit by Thermo Fisher and quality was checked by using Agilent Tapestation. The indexed cDNA libraries were normalized and combined, and the pools were sequenced on the Illumina Nextseq 2000 P3 100 cycles kit sequencing run, generating 58 base paired end reads with dual index 10+10 base pairs.

### Data Analysis

Continuous biological data are reported as mean ± standard deviation.In the biological assumption of normal distributions, histological and biochemical findings were compared by ordinary Two-way ANOVA using GraphPad Prism 8.0 to assess effects of experimental group, strain and interaction. No post-tests between individual columns were performed.

RNA sequencing data underwent basecalling and demultiplexing using CASAVA software with default settings generating Fastq files for further downstream mapping and analysis. Reads were aligned to GRCm38 from Ensembl using STAR (v2.7.9a). Counts for each gene were estimated using featureCounts (v1.5.1). DESeq2 (v.1.42.0) was used for sample group comparisons, generating log2 fold changes, Wald test p-values and p-values adjusted for multiple testing (Benjamini-Hochberg method). GSEA was performed using the fgsea package.^32^ The reference gene sets were obtained from the molecular signatures Database (MSigDB) using the msigdbr package.^33–35^ Specifically from the CP sub-collection of the curated gene set, containing genes from Reactome^36^, WikiPathways^37^ and Biocarta. For each contrast genes were ranked according to a signed significance score (ranking score = sign(log_2_ fold change) x −log_10_(p-value)). This ranking gives positive values to upregulated genes and negative values to downregulated genes, while giving more weight to genes with smaller p-value. Pathways enrichment was then performed with fgsea with standard scoring, including pathways with a size between 10 and 500 genes. Pathways with and adjusted p-value below 0.05 were considered statistically significant. For visualization the package ggplot2^38^ was used. The heatmap was generated by ranking the absolute normalized enrichment score (NES) of all significant pathways and displaying the top 40.

**Figure 1.**
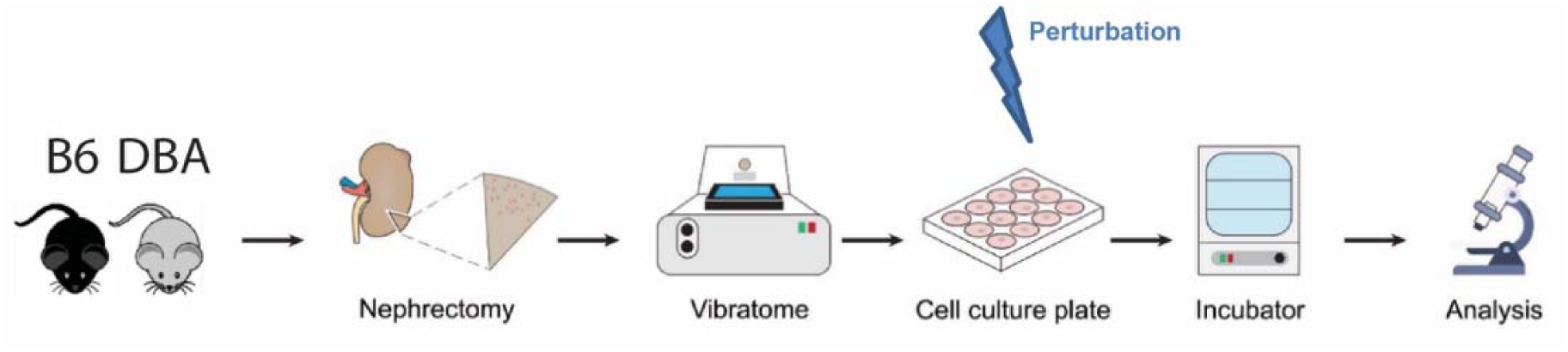
Overview for methodological procedure and analyses: Kidneys were explanted from the mice. A total of 6 animals was used per strain. The kidneys were sectioned to precision cut kidney slices and underwent different perturbation protocols. Experimental readouts included histopathological analysis, TUNEL staining and bulk RNA sequencing.

## Results

Initially, histological analyses were conducted to assess morphological kidney damage in both strains. Overall, a significant difference between the experimental groups was detected (all p < 0.003, 35.6-68.5% of variance explained), whereas no significant main effect of the strain was detected for any morphological parameters (all p > 0.05). Figure 2 summarizes the detailed results for the six morphological parameters evaluated across both mouse strains and all perturbation protocols and corresponding controls. Representative tissue samples are provided in Supplementary Figure 1. The normoxia group was associated with the lowest degree of histopathological tissue injury in both strains, demonstrating significantly less damage compared to both hyperoxia and control groups. Across all morphological parameters, there was a trend towards greater histopathological damage in the DBA/2Jstrain, particularly following 24-hour tunicamycin exposure. No systematic differences were observed between individual perturbation conditions and their respective controls; however, a trend toward increased histopathological tissue damage was noted with prolonged experimental duration. Together, these findings suggest that the extent of histopathological kidney injury was mainly driven by experimental protocol and exposure duration.

**Figure 2.**
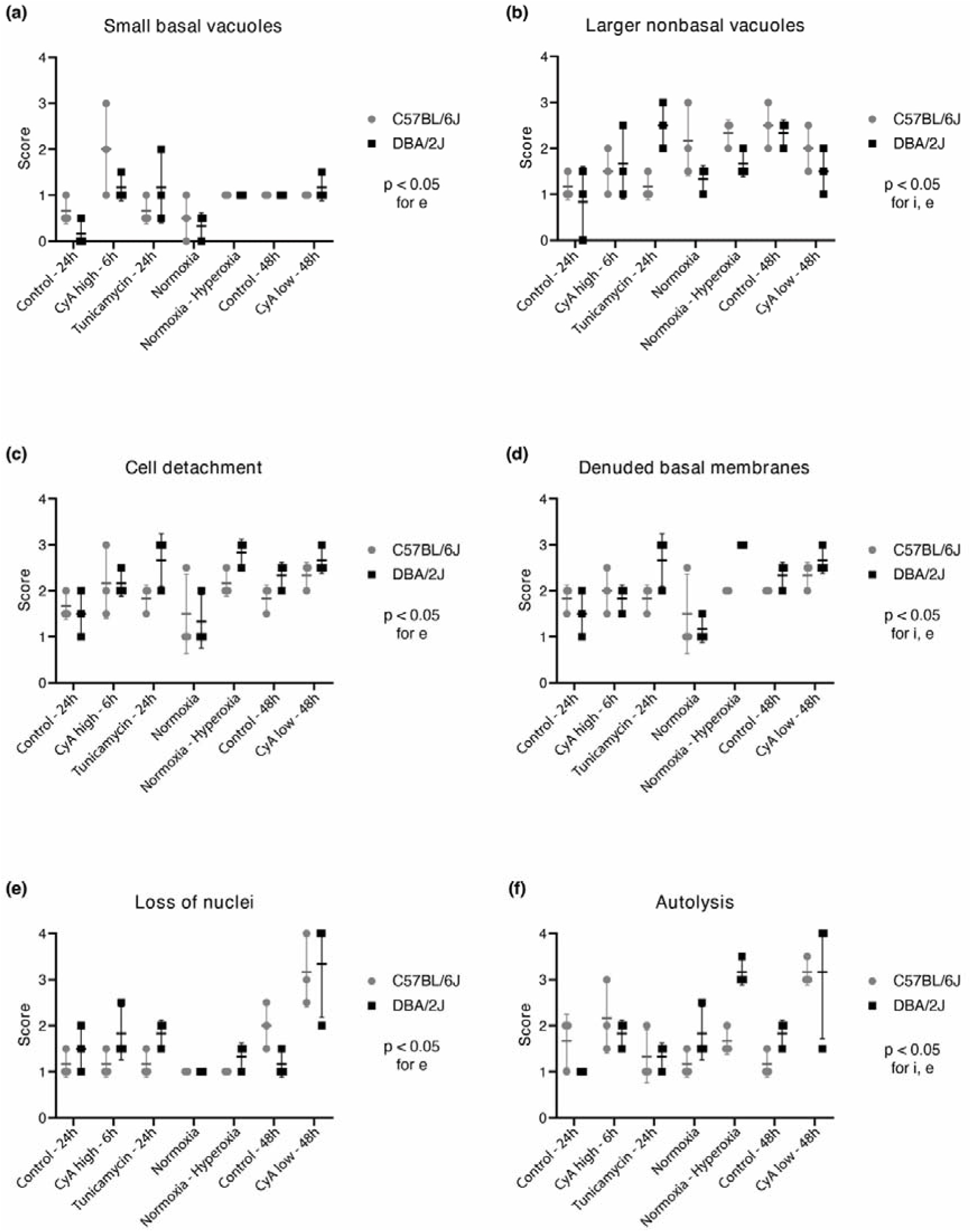
Histopathological assessment of kidney injury: All diagrams show the two-way ANOVA for DBA/2J and C57BL/6J mice for experimental conditions (Cyclosporine A high dose 80 nM 6h, cyclosporine A low dose 20 nM 24h, Tunicamycin 1 µM 24h, Normoxia and Hyperoxia) and the respective controls (24h and 48h). Score definition is based on the extend of tissue affected. Scores indicate 0 = no lesions, 1 = 1-25%, 2 = 26-50%, 3 = 51-75% 4 = 75-100%, i = interaction, e = experimental group, s = strain. (a) shows small basal vacuoles, (b) shows larger nonbasal vacuoles, (c) shows cell detachment, (d) shows denuded basal membranes, (e) shows loss of nuclei and (f) shows autolysis. N = 3 independent samples analysed per condition, per strain.

Subsequently, we aimed to evaluate global metabolic tissue activity by quantifying ATP levels in both strains across all experimental protocols. ATP concentrations differed significantly between the experimental groups (p < 0.0001, 85.2% of variance explained). The DBA/2J strain exhibited consistently lower ATP levels across all conditions (p = 0.0005, explaining 4.3% of variance), with the most pronounced reduction following 6 hours of exposure to high-dose cyclosporine A. Notably, normoxia conditions were associated with the lowest ATP levels in both strains. Detailed results are presented in Figure 3.

**Figure 3.**
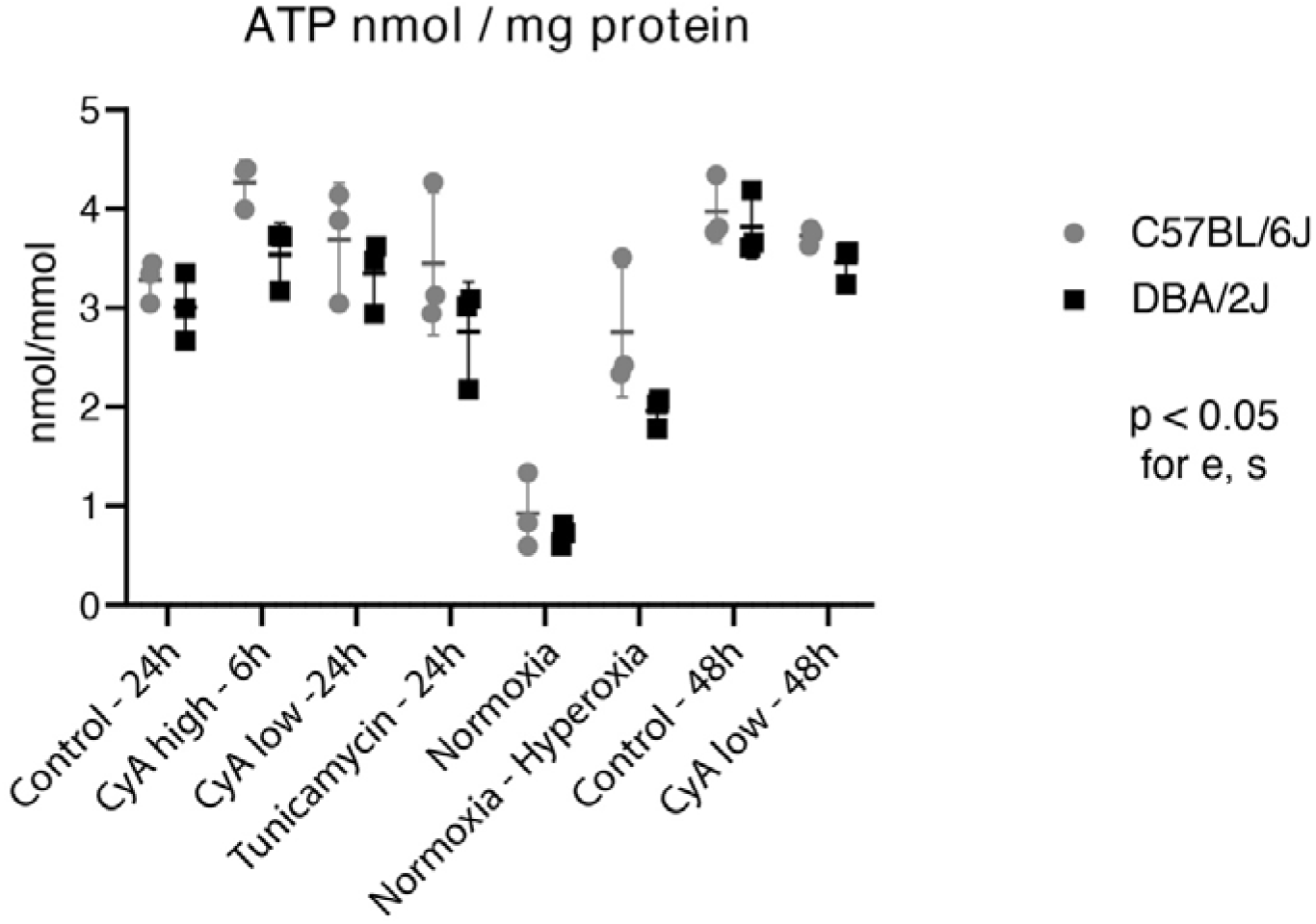
ATP depletion: Two-way ANOVA of ATP levels for both strains DBA/2J and C57BL/6J for experimental conditions (Cyclosporine A high dose 80nM 6h, Cyclosporine A low dose 20nM 24h and 48h, Tunicamycin 1 µM 24h, Normoxia and Hyperoxia) and the corresponding controls (24h and 48h). I = interaction, e = experimental group, s = strain.

To determine whether structural damage and impaired cellular energy metabolism translated into increased cell death, TUNEL-staining was performed to quantify apoptosis. Significant differences in apoptotic cell percentages were observed across experimental groups (p < 0.0001, 62% of variance explained), and additionally a significant interaction between experimental groups and strain was identified (p = 0.006, explaining 15.7% of variance). Figure 4 presents the detailed results and Supplementary Figure 2 shows representative tissue samples. Both normoxia and hyperoxia groups were associated with the lowest proportions of apoptotic cells in both strains. No consistent differences between individual perturbation and their corresponding control were observed; however a trend toward increased apoptotic cell proportion with prolonged experimental duration was noted.

**Figure 4.**
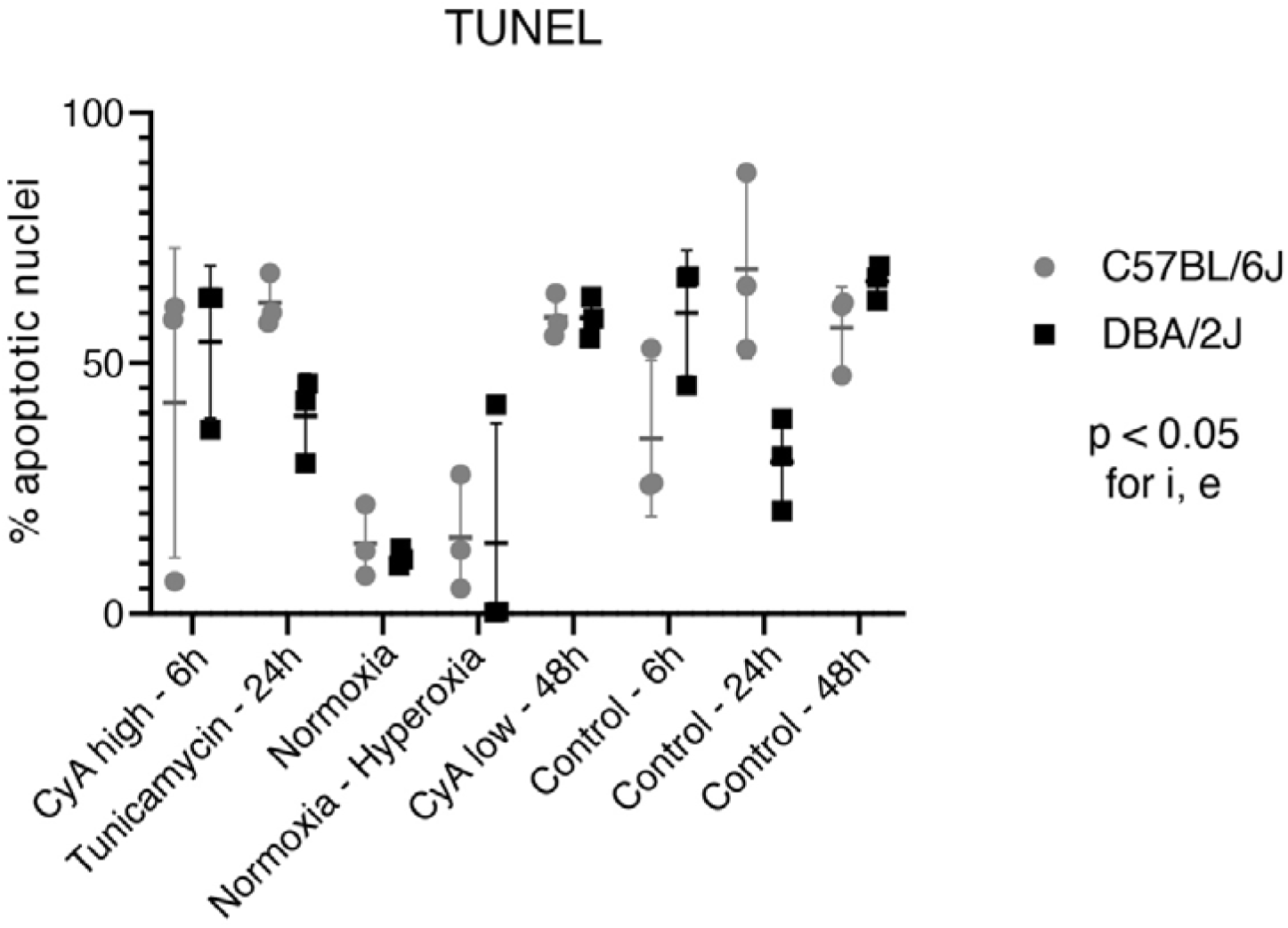
TUNEL-Staining: Two-way ANOVA of percentage of apoptotic nuclei for both strains DBA/2J and C57BL/6J for experimental conditions (Cyclosporine A high dose 80nM 6h, Cyclosporine A low dose 20 nM 48h, Tunicamycin 1 µM 24h, Normoxia and Hyperoxia) and corresponding controls (6h, 24h, 48h). I = interaction, e = experimental group, s = strain.

Finally, RNA sequencing was performed to characterize the underlying transcriptomic response to perturbation. Overall, DBA/2J mice exhibited a stronger response to toxin exposure compared to C57BL/6J mice. First, we performed a principal component analysis, which revealed four different clusters per strain (Figure 5). A normoxia cluster, a hyperoxia cluster, a 6-hour cluster and a 24-hour cluster, with toxin-treated samples and their corresponding controls grouping together within the time-point cluster. The first two principal components accounted for 78% of the total variance. Experimental group explained most of the variance. Individual differentially expressed genes between groups and strains are reported in Supplemental Table 1. Gene set enrichment analysis was subsequently performed to identify pathways differentially regulated between experimental groups and strains. Pathways with the highest normalized enrichment scores (NES) were grouped into different biological categories: extracellular matrix organization, protein metabolism, cell-cycle, aerobic respiration, and metabolism of RNA. Detailed results are presented in Figure 6; Supplementary Figure 3 shows the total number of significant pathways per condition and strain.

**Figure 5.**
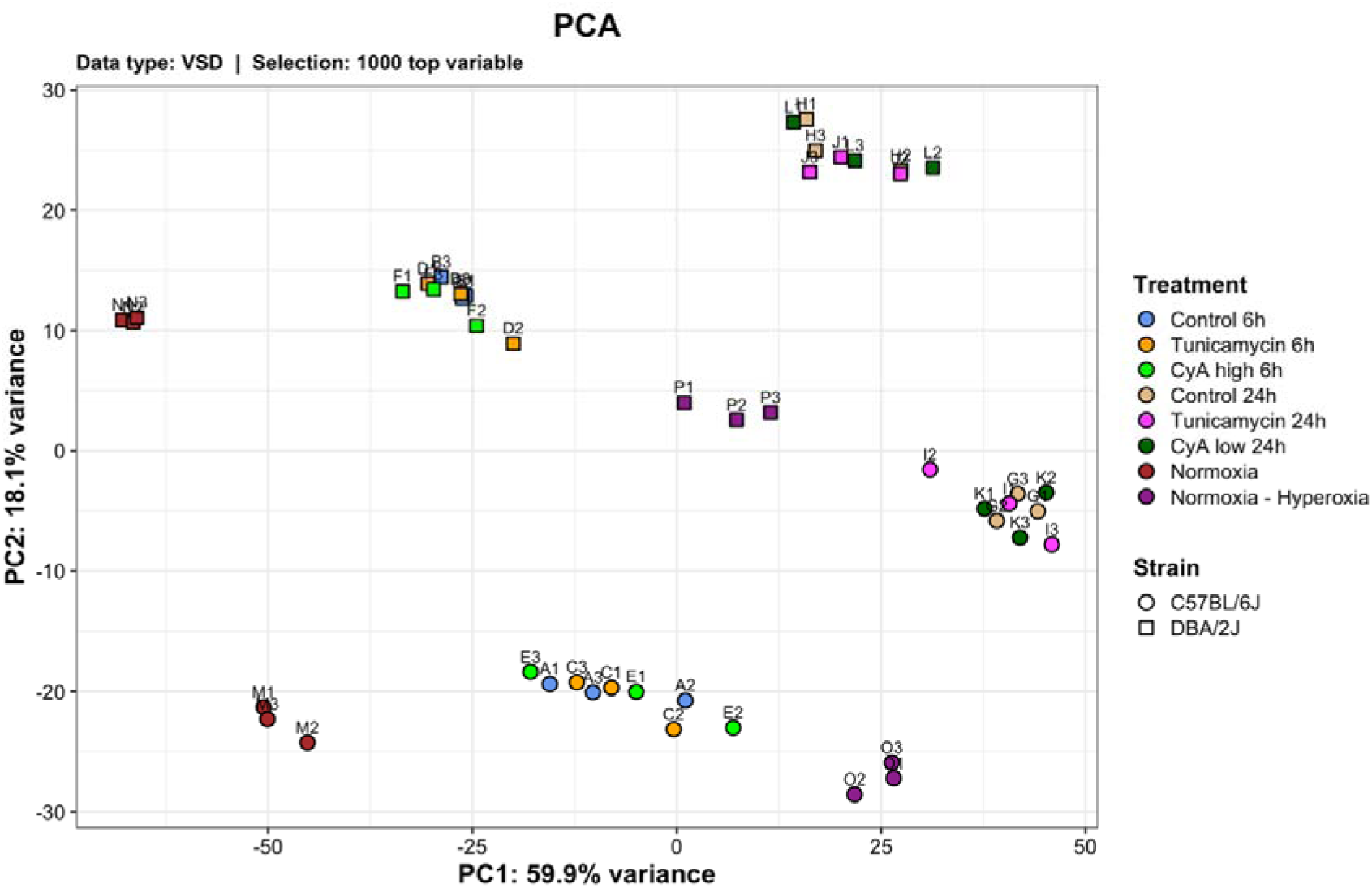
Principal Components Analysis. (PCA) of gene expression data, showing clear clustering of samples for strain and experimental group. The first two principal components (PC1: 59.9% and PC2: 18.1%) account for most of the variance. PC1 primary reflects the effect of the treatment, whereas PC2 depicts variation associated with the effect of the strain.

**Figure 6.**
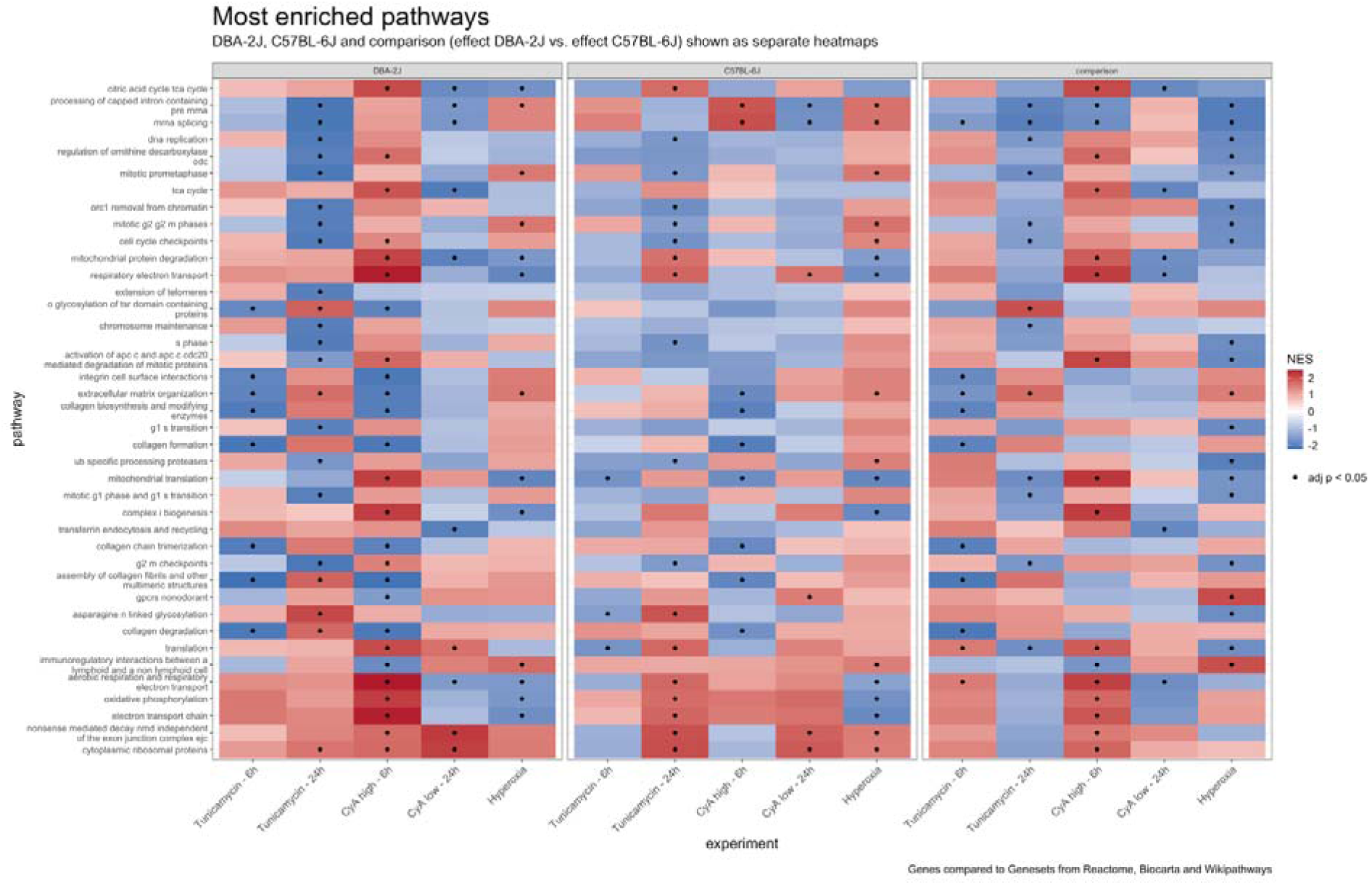
Gene set enrichment analysis: Heatmap of most enriched pathways from the selection of Reactome, Biocarta and Wikipathways. Left and middle panel show the comparison of experimental groups to the corresponding control and hyperoxia compared to normoxia per strain. The left panel for DBA/2J and in the middle panel for C57BL/6J. The right panel shows the comparison between DBA/2J and C7BL-6J. Significant Pathways were sorted by absolute NES and the top 30 pathways displayed.

Extracellular Matrix (ECM) organization pathways were downregulated after 6-hour exposure to tunicamycin and high-dose cyclosporine A in both strains, with significantly stronger downregulation in DBA/2J mice. Following hyperoxia ECM organization pathways were upregulated in both strains. Protein metabolism pathways included predominantly translational and posttranslational processes. Following 6-hour exposure to tunicamycin and high-dose cyclosporine A, most pathways were upregulated in DBA/2J mice and downregulated in C57BL/6J mice. Direct comparison therefore revealed significantly stronger enrichment of these pathways in DBA/2J mice. Cell-cycle pathways were downregulated after 24-hour tunicamycin exposure in both strains, but in comparison significantly less enriched in DBA/2J mice. Following hyperoxia, these pathways were significantly upregulated in C57BL/6J mice, while no clear trend was observed in DBA/2J mice. Cell-cycle pathway enrichment was significantly stronger in C57BL/6J mice. Pathways associated with aerobic respiration were upregulated after 6-hour exposure in DBA/2J mice and significantly more enriched than in C57BL/6J mice, particularly following 6-hour high-dose cyclosporine A exposure. After 24-hour exposure to tunicamycin and low-dose cyclosporine A these pathways were upregulated in both strains, except for 24-hour cyclosporine A exposure in DBA/2J mice. Direct comparison revealed stronger enrichment in C57BL/6J mice at the 24-hour time point. Taken together, these results suggest that transcriptomic alterations were largely driven by the experimental protocol and duration, while strain-specific characteristics further modulated the response, with DBA/2J mice showing a stronger response to early toxin perturbation, with less ECM formation and cell-cycle activity and more aerobic respiration and protein metabolism.

## Discussion

In this study we set out to investigate whether to two founder strains of the BXD mouse consortium, DBA/2J and C57BL/6J, differ in their response to nephrotoxic perturbation using precision cut kidney slices. We found that the main source of variation was the time of the experimental duration. Nevertheless, we observed subtle differences between the two strains, with DBA/2J mice showing a stronger response to nephrotoxic stress. This supports previous evidence that DBA/2J mice are more susceptible for kidney damage, for example in diabetic nephropathy^39^ and adenine-induced kidney injury in unpublished data from Karolinska institute in Stockholm. The observed differences suggest that the BXD-Consortium shows different susceptibility to nephrotoxins. DBA/2J and C57BL/6J do not necessarily represent the most extreme phenotypes within the full BXD population.^40^ Therefore, the differences detected between these founder strains provide sufficient rationale for further investigation in the broader BXD resource.

The data obtained from bulk RNA sequencing underlines the biological relevance of the observed injury response. Gene set enrichment analysis showed a downregulation of extracellular matrix organization and cell-cycle pathways. Previous studies have associated these pathways with impaired tissue maintenance, reduced regenerative capacity and early injury associated remodelling processes during acute kidney injury.^41–43^ Although the transcriptional differences between the two strains were modest, the stronger downregulation of these pathways in DBA/2J mice aligns with their increased susceptibility to drug-induced kidney injury. Together, the transcriptional changes suggest that the PCKS model captures molecular features that are consistent with early injury response in acute kidney injury and may therefore provide a suitable model for studying genetically driven differences in renal stress responses.

Precision-cut tissue slices have been widely used in liver toxicity research, including studies on idiosyncratic drug toxicity, cholestatic liver damage, and drug-induced apoptosis^44–46^, supporting their value as ex vivo model for investigating drug toxicity. The validity of PCKS as an ex vivo model is further supported by studies using renal cortical tissue slices. Bavere et al.^47^ showed that PCKS can be used to investigate drug-induced renal injury, while Arakawa et al.^48^ demonstrated that rat kidney slices can be used to detect platinum-induced kidney injury. Together these studies support the use of PCKS as physiologically relevant ex vivo model for nephrotoxicity research.

PCKS have several advantages when compared with other acute kidney injury models. In vitro cell cultures are traditionally used to look at a single cell type, with the kidney being one of the most complex organs it is difficult to model the process of a AKI, as many cell type types are involved in the pathogenesis.^49^ In contrast to cell cultures, PCKS preserve tissue architecture and maintain all cell-cell contacts, allowing the investigation of drug efficacy, toxicity and metabolism in a more physiologically relevant environment. Many previous studies have shown that rodent models show clear limitations in capturing human pathophysiology in acute kidney injury.^50–53^ In addition preparing PCKS is relatively efficient, as multiple slices can be obtained from a small amount of tissue and they can be prepared from both animal and human renal tissue.^22^ Unlike in vivo models, PCKS allow the investigation of early tissue response under highly controlled experimental conditions in a more organ specific context. PCKS therefore offers a chance to investigate AKI in slices obtained from human kidneys for example after explanation or from biopsies a controlled environment. These characteristics make PCKS particularly suitable for studying early mechanisms of drug-induced kidney injury.

Nevertheless, PCKS also has limitations, some of which became apparent in the present study. Maintaining slice viability remains challenging, limiting the experimental duration to approximately 48 hours. This time frame may be too short to detect pronounced histological damage, which could explain why no histological differences between strains could be observed, while ATP levels, apoptosis and transcriptomes showed subtle differences between the two strains. Furthermore, experimental duration appeared to be the major source of variation, suggesting that experimental condition themselves cause tissue damage. This highlights the importance of strict standardization of experimental conditions, incubation times and sample handling during larger genomic studies based on PCKS to ensure comparable and reproductible results. Thus, our findings suggest that PCKS are particularly useful for detecting early functional and molecular responses to nephrotoxic stress. Additional limitations of this study include the relatively small sample size and small number of strains, as well as the use of only male mice. Therefore, potential sex-specific differences in nephrotoxin susceptibility could not be assessed.

Previous studies have mainly focused on modelling nephrotoxicity, fibrosis or drug-induced injury in general.^22,27,28,47^ Our study shows that PCKS can be used to investigate genetically determined susceptibility to nephrotoxic injury in the BXD founder strains. This provides the foundation for extending the PCKS approach to the full BXD population. Further studies should therefore include additional BXD strains to determine whether the observed differences between DBA/2J and C57BL/6J mice translate to the broader genetic variation within the consortium, and perform heritability estimation and ultimately QTL mapping to identify novel genetic loci and specific candidate genes associated with susceptibility to drug-induced kidney injury.

In addition, future research should investigate other clinically relevant nephrotoxic drugs, as current studies are still largely focused on genetic differences in platinum-mediated kidney injury. Technical optimizations of the PCKS culture system might also improve model performance. Ultimately, preparing PCKS from human renal tissue may offer a translational perspective, as this approach could be used to better understand patient-specific susceptibility to nephrotoxic drugs prediction.g. in the case of renal cancer where non-tumore kidney tissue from nephrectomies is routinely available.

## Acknowledgements

The authors would like to thank BEA, the core facility for Bioinformatics and Expression Analysis, which is supported by the board of research at the Karolinska Institutet and the research committee at the Karolinska hospital. The authors also acknowledge staff of the FENO Morphological Phenotype Analysis Core for histology services.

## Author contributions

JA performed computational analyses and wrote the manuscript. NP performed wet-lab experiments. MB performed animal experiments. HO and JP provided research infrastructure and critical reagents. MBM designed and supervised the project. All authors critically reviewed the manuscript and agreed to publication.

## Accession codes

The murine bulk RNA-Seq datasets generated within this study are accessible at Gene Expression Omnibus under the accession number GSE334849.

## References

1. Mehta RL, Cerdá J, Burdmann EA, et al. International Society of Nephrology’s 0by25 initiative for acute kidney injury (zero preventable deaths by 2025): a human rights case for nephrology. Lancet. 2015;385(9987):2616–2643. doi:10.1016/S0140-6736(15)60126-X

2. Nash K, Hafeez A, Hou S. Hospital-acquired renal insufficiency. American Journal of Kidney Diseases. 2002;39(5):930–936. doi:10.1053/ajkd.2002.32766

3. Ortega-Loubon C, Martínez-Paz P, García-Morán E, et al. Genetic Susceptibility to Acute Kidney Injury. J Clin Med. 2021;10(14):3039. doi:10.3390/jcm10143039

4. Zuk A, Bonventre JV. Acute Kidney Injury. Annu Rev Med. 2016;67:293–307. doi:10.1146/annurev-med-050214-013407

5. Kaufman J, Dhakal M, Patel B, Hamburger R. Community-Acquired Acute Renal Failure. American Journal of Kidney Diseases. 1991;17(2):191–198. doi:10.1016/S0272-6386(12)81128-0

6. Chawla LS, Eggers PW, Star RA, Kimmel PL. Acute kidney injury and chronic kidney disease as interconnected syndromes. N Engl J Med. 2014;371(1):58–66. doi:10.1056/NEJMra1214243

7. Ishani A, Xue JL, Himmelfarb J, et al. Acute Kidney Injury Increases Risk of ESRD among Elderly. Journal of the American Society of Nephrology. 2009;20(1):223–228. doi:10.1681/ASN.2007080837

8. Gandhi TK, Burstin HR, Cook EF, et al. Drug complications in outpatients. J Gen Intern Med. 2000;15(3):149–154. doi:10.1046/j.1525-1497.2000.04199.x

9. Bellomo R. The epidemiology of acute renal failure: 1975 versus 2005: Current Opinion in Critical Care. 2006;12(6):557–560. doi:10.1097/01.ccx.0000247443.86628.68

10. Perazella MA, Rosner MH. Drug-Induced Acute Kidney Injury. Clin J Am Soc Nephrol. 2022;17(8):1220–1233. doi:10.2215/CJN.11290821

11. Naughton CA. Drug-induced nephrotoxicity. Am Fam Physician. 2008;78(6):743–750.

12. Klumpers MJ, Witte WD, Gattuso G, et al. Genome-Wide Analyses of Nephrotoxicity in Platinum-Treated Cancer Patients Identify Association with Genetic Variant in RBMS3 and Acute Kidney Injury. J Pers Med. 2022;12(6):892. doi:10.3390/jpm12060892

13. Andreux PA, Williams EG, Koutnikova H, et al. Systems genetics of metabolism: the use of the BXD murine reference panel for multiscalar integration of traits. Cell. 2012;150(6):1287–1299. doi:10.1016/j.cell.2012.08.012

14. Wang X, Pandey AK, Mulligan MK, et al. Joint mouse-human phenome-wide association to test gene function and disease risk. Nat Commun. 2016;7:10464. doi:10.1038/ncomms10464

15. Peirce JL, Lu L, Gu J, Silver LM, Williams RW. A new set of BXD recombinant inbred lines from advanced intercross populations in mice. BMC Genet. 2004;5:7. doi:10.1186/1471-2156-5-7

16. Wang X, Agarwala R, Capra JA, et al. High-throughput sequencing of the DBA/2J mouse genome. BMC Bioinformatics. 2010;11(S4):O7. doi:10.1186/1471-2105-11-S4-O7

17. Mouse Genome Sequencing Consortium, Waterston RH, Lindblad-Toh K, et al. Initial sequencing and comparative analysis of the mouse genome. Nature. 2002;420(6915):520–562. doi:10.1038/nature01262

18. Clee SM, Yandell BS, Schueler KM, et al. Positional cloning of Sorcs1, a type 2 diabetes quantitative trait locus. Nat Genet. 2006;38(6):688–693. doi:10.1038/ng1796

19. Wang X, Ria M, Kelmenson PM, et al. Positional identification of TNFSF4, encoding OX40 ligand, as a gene that influences atherosclerosis susceptibility. Nat Genet. 2005;37(4):365–372. doi:10.1038/ng1524

20. Klein RF, Allard J, Avnur Z, et al. Regulation of bone mass in mice by the lipoxygenase gene Alox15. Science. 2004;303(5655):229–232. doi:10.1126/science.1090985

21. Wu Y, Williams EG, Dubuis S, et al. Multilayered genetic and omics dissection of mitochondrial activity in a mouse reference population. Cell. 2014;158(6):1415–1430. doi:10.1016/j.cell.2014.07.039

22. Poosti F, Pham BT, Oosterhuis D, et al. Precision-cut kidney slices (PCKS) to study development of renal fibrosis and efficacy of drug targeting *ex vivo*. Disease Models & Mechanisms. Published online January 1, 2015:dmm.020172. doi:10.1242/dmm.020172

23. Stribos EGD, Seelen MA, van Goor H, Olinga P, Mutsaers HAM. Murine Precision-Cut Kidney Slices as an ex vivo Model to Evaluate the Role of Transforming Growth Factor-β1 Signaling in the Onset of Renal Fibrosis. Front Physiol. 2017;8:1026. doi:10.3389/fphys.2017.01026

24. Klinkhammer BM, Goldschmeding R, Floege J, Boor P. Treatment of Renal Fibrosis—Turning Challenges into Opportunities. Advances in Chronic Kidney Disease. 2017;24(2):117–129. doi:10.1053/j.ackd.2016.11.002

25. Stribos EGD, Hillebrands JL, Olinga P, Mutsaers HAM. Renal fibrosis in precision-cut kidney slices. European Journal of Pharmacology. 2016;790:57–61. doi:10.1016/j.ejphar.2016.06.057

26. Charrin E, Dabaghie D, Sen I, et al. Soluble Klotho protects against glomerular injury through regulation of ER stress response. Commun Biol. 2023;6(1):208. doi:10.1038/s42003-023-04563-1

27. Moor M, Nordström J, Burmakin M, et al. Comparative analysis of kidney transplantation modeled using precision-cut kidney slices and kidney transplantation in pigs. Nephrology Dialysis Transplantation. 2024;39(Supplement_1):gfae069-0953-1945. doi:10.1093/ndt/gfae069.953

28. Bigaeva E, Puerta Cavanzo N, Stribos EGD, et al. Predictive Value of Precision-Cut Kidney Slices as an Ex Vivo Screening Platform for Therapeutics in Human Renal Fibrosis. Pharmaceutics. 2020;12(5):459. doi:10.3390/pharmaceutics12050459

29. Hubrecht RC, Carter E. The 3Rs and Humane Experimental Technique: Implementing Change. Animals (Basel). 2019;9(10):754. doi:10.3390/ani9100754

30. Bankhead P, Loughrey MB, Fernández JA, et al. QuPath: Open source software for digital pathology image analysis. Sci Rep. 2017;7(1):16878. doi:10.1038/s41598-017-17204-5

31. Love MI, Huber W, Anders S. Moderated estimation of fold change and dispersion for RNA-seq data with DESeq2. Genome Biol. 2014;15(12):550. doi:10.1186/s13059-014-0550-8

32. Korotkevich G, Sukhov V, Budin N, Shpak B, Artyomov MN, Sergushichev A. Fast gene set enrichment analysis. Bioinformatics. Preprint posted online June 20, 2016. doi:10.1101/060012

33. Subramanian A, Tamayo P, Mootha VK, et al. Gene set enrichment analysis: a knowledge-based approach for interpreting genome-wide expression profiles. Proc Natl Acad Sci U S A. 2005;102(43):15545–15550. doi:10.1073/pnas.0506580102

34. Liberzon A, Subramanian A, Pinchback R, Thorvaldsdóttir H, Tamayo P, Mesirov JP. Molecular signatures database (MSigDB) 3.0. Bioinformatics. 2011;27(12):1739–1740. doi:10.1093/bioinformatics/btr260

35. Castanza AS, Recla JM, Eby D, Thorvaldsdóttir H, Bult CJ, Mesirov JP. Extending support for mouse data in the Molecular Signatures Database (MSigDB). Nat Methods. 2023;20(11):1619–1620. doi:10.1038/s41592-023-02014-7

36. Ragueneau E, Gong C, Sinquin P, et al. The Reactome Knowledgebase 2026. Nucleic Acids Res. 2026;54(D1):D673–D681. doi:10.1093/nar/gkaf1223

37. Agrawal A, Balcı H, Hanspers K, et al. WikiPathways 2024: next generation pathway database. Nucleic Acids Research. 2024;52(D1):D679–D689. doi:10.1093/nar/gkad960

38. Wickham H. Ggplot2: Elegant Graphics for Data Analysis. Springer-Verlag New York; 2016. https://ggplot2.tidyverse.org

39. Østergaard MV, Pinto V, Stevenson K, Worm J, Fink LN, Coward RJM. DBA/2J2J *db/db* mice are susceptible to early albuminuria and glomerulosclerosis that correlate with systemic insulin resistance. American Journal of Physiology-Renal Physiology. 2017;312(2):F312–F321. doi:10.1152/ajprenal.00451.2016

40. Wang Q, Qi H, Wu Y, et al. Genetic susceptibility to diabetic kidney disease is linked to promoter variants of XOR. Nat Metab. 2023;5(4):607–625. doi:10.1038/s42255-023-00776-0

41. Yang L, Besschetnova TY, Brooks CR, Shah JV, Bonventre JV. Epithelial cell cycle arrest in G2/M mediates kidney fibrosis after injury. Nat Med. 2010;16(5):535–543, 1p following 143. doi:10.1038/nm.2144

42. Janosevic D, De Luca T, Melo Ferreira R, et al. miRNA and mRNA Signatures in Human Acute Kidney Injury Tissue. Am J Pathol. 2025;195(1):102–114. doi:10.1016/j.ajpath.2024.08.013

43. Baker ML, Cantley LG. Adding insult to injury: the spectrum of tubulointerstitial responses in acute kidney injury. J Clin Invest. 2025;135(6):e188358. doi:10.1172/JCI188358

44. Vatakuti S, Olinga P, Pennings JLA, Groothuis GMM. Validation of precision-cut liver slices to study drug-induced cholestasis: a transcriptomics approach. Arch Toxicol. 2017;91(3):1401–1412. doi:10.1007/s00204-016-1778-8

45. Martinez SM, Bradford BU, Soldatow VY, et al. Evaluation of an in vitro toxicogenetic mouse model for hepatotoxicity. Toxicol Appl Pharmacol. 2010;249(3):208–216. doi:10.1016/j.taap.2010.09.012

46. Hadi M, Chen Y, Starokozhko V, Merema MT, Groothuis GMM. Mouse precision-cut liver slices as an ex vivo model to study idiosyncratic drug-induced liver injury. Chem Res Toxicol. 2012;25(9):1938–1947. doi:10.1021/tx300248j

47. Baverel G, Knouzy B, Gauthier C, et al. Use of precision-cut renal cortical slices in nephrotoxicity studies. Xenobiotica. 2013;43(1):54–62. doi:10.3109/00498254.2012.725142

48. Arakawa H, Nagao Y, Nedachi S, Shirasaka Y, Tamai I. Evaluation of Platinum Anticancer Drug-Induced Kidney Injury in Primary Culture of Rat Kidney Tissue Slices by Using Gas-Permeable Plates. Biol Pharm Bull. 2022;45(3):316–322. doi:10.1248/bpb.b21-00875

49. Kishi S, Matsumoto T, Ichimura T, Brooks CR. Human reconstructed kidney models. In Vitro Cell Dev Biol Anim. 2021;57(2):133–147. doi:10.1007/s11626-021-00548-8

50. Lindström NO, McMahon JA, Guo J, et al. Conserved and Divergent Features of Human and Mouse Kidney Organogenesis. J Am Soc Nephrol. 2018;29(3):785–805. doi:10.1681/ASN.2017080887

51. Miyoshi T, Hiratsuka K, Saiz EG, Morizane R. Kidney organoids in translational medicine: Disease modeling and regenerative medicine. Dev Dyn. 2020;249(1):34–45. doi:10.1002/dvdy.22

52. Odom DT, Dowell RD, Jacobsen ES, et al. Tissue-specific transcriptional regulation has diverged significantly between human and mouse. Nat Genet. 2007;39(6):730–732. doi:10.1038/ng2047

53. de Caestecker M, Humphreys BD, Liu KD, et al. Bridging Translation by Improving Preclinical Study Design in AKI. J Am Soc Nephrol. 2015;26(12):2905–2916. doi:10.1681/ASN.2015070832

