## Supplemental Figures for "Population genetics meets precision-cut kidney slices: Nephrotoxicity modelled ex vivo in the founder strains of the BXD mouse consortium"

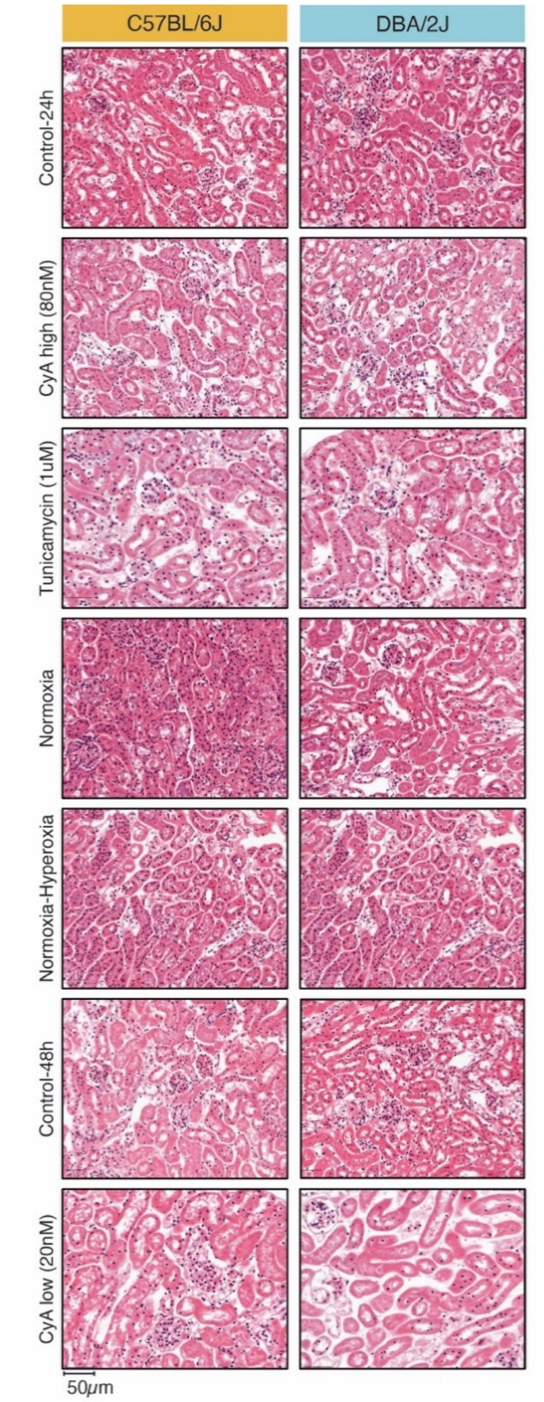


**Supplemental Figure 1**

Representative H&E-stained tissue sections. Example of the haematoxylin-eosin-stained tissue used for scoring. Left panel shows C57BL/6J, right panel shows DBA/2J.

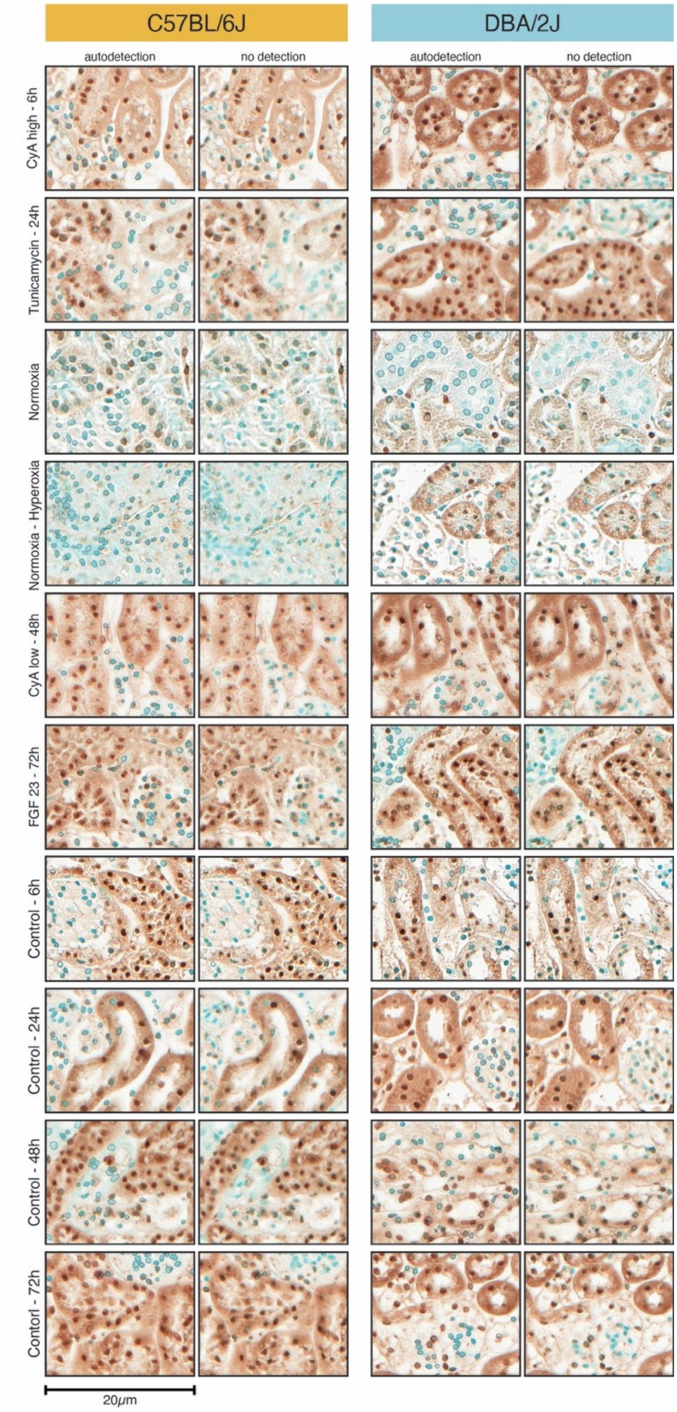


**Supplemental Figure 2**

Representative TUNEL stained tissue sections. Left panel shows C57BL/6J, right panel shows DBA/2J. Left columns in each group show the Autodetection by QuPath; red = positive nuclei, blue = negative nuclei, right columns show the tissue section without detections.

**Summary of significant pathways**

| | Strain | n pathways | n upregulated | n downregulated | mean NES upregulated | mean NES downregulated |
|---|---|---|---|---|---|---|
| **Tunicamycin - 6h** | DBA-2J | 15 | 0 | 15 | NA | –1.97 |
| **Tunicamycin - 6h** | C57BL-6J | 19 | 0 | 19 | NA | –1.66 |
| **Tunicamycin - 6h** | comparison | 19 | 9 | 10 | 1.69 | –1.89 |
| **CyA high - 6h** | DBA-2J | 121 | 45 | 76 | 1.81 | –1.71 |
| **CyA high - 6h** | C57BL-6J | 39 | 27 | 12 | 1.68 | –1.85 |
| **CyA high - 6h** | comparison | 113 | 46 | 67 | 1.79 | –1.66 |
| **Tunicamycin - 24h** | DBA-2J | 174 | 22 | 152 | 1.76 | –1.77 |
| **Tunicamycin - 24h** | C57BL-6J | 66 | 22 | 44 | 1.78 | –1.66 |
| **Tunicamycin - 24h** | comparison | 39 | 11 | 28 | 1.77 | –1.69 |
| **CyA low - 24h** | DBA-2J | 43 | 6 | 37 | 1.94 | –1.72 |
| **CyA low - 24h** | C57BL-6J | 10 | 5 | 5 | 1.66 | –1.65 |
| **CyA low - 24h** | comparison | 8 | 1 | 7 | 1.83 | –1.86 |
| **Hyperoxia** | DBA-2J | 123 | 87 | 36 | 1.61 | –1.63 |
| **Hyperoxia** | C57BL-6J | 228 | 184 | 44 | 1.53 | –1.66 |
| **Hyperoxia** | comparison | 209 | 41 | 168 | 1.70 | –1.71 |

**Supplemental Figure 3.**

Summary of significant pathways per condition per strain. Number of significant pathways and mean NES for upregulated and downregulated pathways across the experimental groups per strain. Experimental groups were compared to the corresponding control, Hyperoxia was compared with normoxia and in the comparison group DBA/2J was compared to C57BL/6J.
